# Pixel intensity of wing photos used to predict age of *Anopheles gambiae* caught during the RIMDAMAL II clinical trial

**DOI:** 10.1101/2025.02.27.640698

**Authors:** Greg Pugh, Emmanuel Sougué, Timothy A. Burton, Ryan Yoe, Lyndsey I Gray, Blue Hephaestus, Anthony Somé, Bryce Asay, Sunil Parikh, A. Fabrice Somé, Roch K. Dabiré, Brian D. Foy

## Abstract

Mosquito age-grading is important for evaluating mosquito control efforts and estimating pathogen transmission risk. We previously developed a simple, low-cost, and high-throughput method to age-grade mosquitoes by computing the pixel intensity (PI) of wing photos, which reflects wing scale loss over time. Here the technique was refined and used to understand wild *Anopheles gambiae* population structures from the RIMDAMAL II clinical trial. Wing photos from lab-reared *An. gambiae* had narrower PI ranges compared to wild *An. gambiae* s.l., but the PI distributions reflected wild population structures where most have lower PI values and very few high PI values. A model was then fitted to samples from lab mosquitoes of known ages and used to interpolate unknown ages from the wild populations (median age of 4.96 [0.8-14.93] days old). Analyses from the RIMDAMAL II trial indicate that while ivermectin mass drug administrations in the intervention arm may have modestly influenced PI and predicted age relative to controls, distributions of new dual-chemistry bed nets in both arms were associated with a strong effect on PI and predicted age structure. These data suggest this method can be used to rapidly age-grade wild mosquito populations and infer the efficacy of vector control interventions.

## Introduction

An adult female mosquito’s age is a key factor in its ability to blood feed, lay eggs and transmit pathogens. Anthropophilic species like *Anopheles gambiae* typically obtain their first bloodmeal 3-5 days post-eclosion^1^. The extrinsic incubation period (EIP), defines the time it takes for a vector-borne pathogen ingested in a competent mosquito’s blood meal to infect or penetrate the mosquito’s tissues, undergo development (in the case of *Plasmodium* parasites or filarial worms) and disseminate to tissues from where it can be transmitted to the next vertebrate host (e.g. salivary glands or thoracic flight muscles). For *Plasmodium falciparum* in African *Anopheles* spp., the EIP is approximately between 9-16 days depending on temperature and species^2,3^, while the typical lifespan of an *Anopheles* mosquito in the wild is roughly 15-20 days^3–5^. Thus, very few mosquitoes in populations ever become infectious and large sample sizes are required to reliably estimate transmission intensity. However, much effort, time and costs can be expended to capture and test for infectious mosquitoes from these large collections. As such, the ability to rapidly and accurately age-grade captured mosquitoes from a population could allow for simple and less resource-intensive estimates of transmission risk and help better evaluate the efficacy of vector control interventions that often occur before periods of high pathogen transmission.

Original mosquito age-grading techniques were qualitative, such as classifying wing degradation, assessing the presence of mites on mosquitoes, or meconium in midguts^6–8^. Eventually evaluating parity by assessing ovary tracheole skeins on dissected ovaries became the gold standard age grading technique, which was further refined and quantified by counting ovariole dilatations from dissected ovaries^9–11^.

However, these techniques can be imprecise, they only provide physiological age rather than calendar age, and it can be difficult and time-consuming to process many mosquitoes per field collection with the dissection techniques required. New age-grading techniques have been investigated, including biochemical, gene and protein profiling, and spectroscopy methods^12–25^. While these methods are quickly reinvigorating mosquito age-grading processes, there remain challenges towards practical implementation in the field. These challenges include the need for expensive equipment which can be difficult to use in a field setting, or shipment of specimens for analysis by this equipment outside of the field site, as well as destructive testing of the samples. Furthermore, these techniques may require recalibration of age models at each field site and/or timepoint the samples are collected, and they can involve advanced technical and statistical expertise to run samples and analyze the data produced^21,26^.

We previously developed a simple, low-cost, high-throughput, and nondestructive method of age-grading lab-reared *Anopheles gambiae* populations by calculating the pixel intensity (PI) of wing photos, which analyzes the intensity (darkness) of the pixels of a mosquito wing photo in grayscale^27^. This method can quantitatively assess wing scale loss over time due to flying because a mosquito’s wing scales are often darker than the underlying membranous wing. In the study, we demonstrated this method’s ability to distinguish the age structure of two mosquito populations reared in different mesocosms, whereby one was regularly treated with blood meals that contained the mosquitocidal drug ivermectin, and the other control mesocosm was only provided regular blood meals. However, it was unclear if this method could be extended to field mosquitoes. Here, we have refined our techniques for PI analysis of wings, applied our lab-reared *An. gambiae* age-model to estimate ages of wild caught *An. gambiae* s.l. captured during a clinical trial conducted in Burkina Faso called RIMDAMAL II (which tested the effect of ivermectin mass drug administrations (MDA) on malaria incidence)^28^, and demonstrate how this technique can infer the efficacy of vector control interventions that occurred during the trial.

## Results

### Total Pixel Intensity vs Mean Pixel Intensity

Our previously published PI analyses calculated the total PI of each wing photo (left– and right-wing photos) per mosquito and used these values to calculate the mean of the total PI per mosquito ^27^. However, this analysis yielded highly variable results of the mean total PI from some mosquitoes caught and processed during the RIMDAMAL II trial, which we determined were due to differences in file sizes, photo resolution, and relative photo brightness. Independently analyzing the mean PI of each wing photo normalized the PI output regardless of file size and other parameters. A mean PI per mosquito was then calculated using the mean PI of each wing, which allowed for normalized comparisons across groups of mosquitoes processed during the trial.

Overall, the prior published analysis of total PI gave ranges from 2.4×10^8^-6.6×10^8^ while our updated analysis of mean PI gave ranges from 127.85-133.08.

### PI Standard Deviation Among Mosquito Wings

We then calculated the standard deviation (SD) of the mean PI between the two wings per mosquito to understand how the variance between wings may be associated with the mean PI and thus potentially influence a mosquito’s predicted age. Among the lab mosquitoes aged in a mesocosm, the mean SD was 0.13 (Range: 0.00-0.76) and among the wild mosquitoes, the mean SD was 0.08 (Range: 0.0004-3.34), however, comparisons of SD of both mosquito populations relative to mean PI or age suggested the SD of the PI between mosquitoes’ wings has only a modest effect on the overall estimated age distribution of the analyzed mosquito population (Figure 1). Overall, among wild mosquitoes where the SD between wings was less than 0.5, the mean PI was 128.51 (Range: 127.82-133.15) while those where the SD between wings was greater than 0.5 had a mean PI of 130.72 (Range: 128.44-133.07), suggesting that as a mosquito ages and loses more scales through more flying, there is a modestly higher likelihood of having one wing that has significantly fewer scales than the other. A similar trend is observed in the lab mosquitoes. Finally, analysis of each pair of wings suggests there is no favoritism between left and right wing. 51.69% (95% 49.42 – 53.96%) of left wings had a higher PI while 48.31% (95% CI: 46.04 – 50.58%) right wings had a higher PI (Supplemental Figure 2).

**Figure 1.**
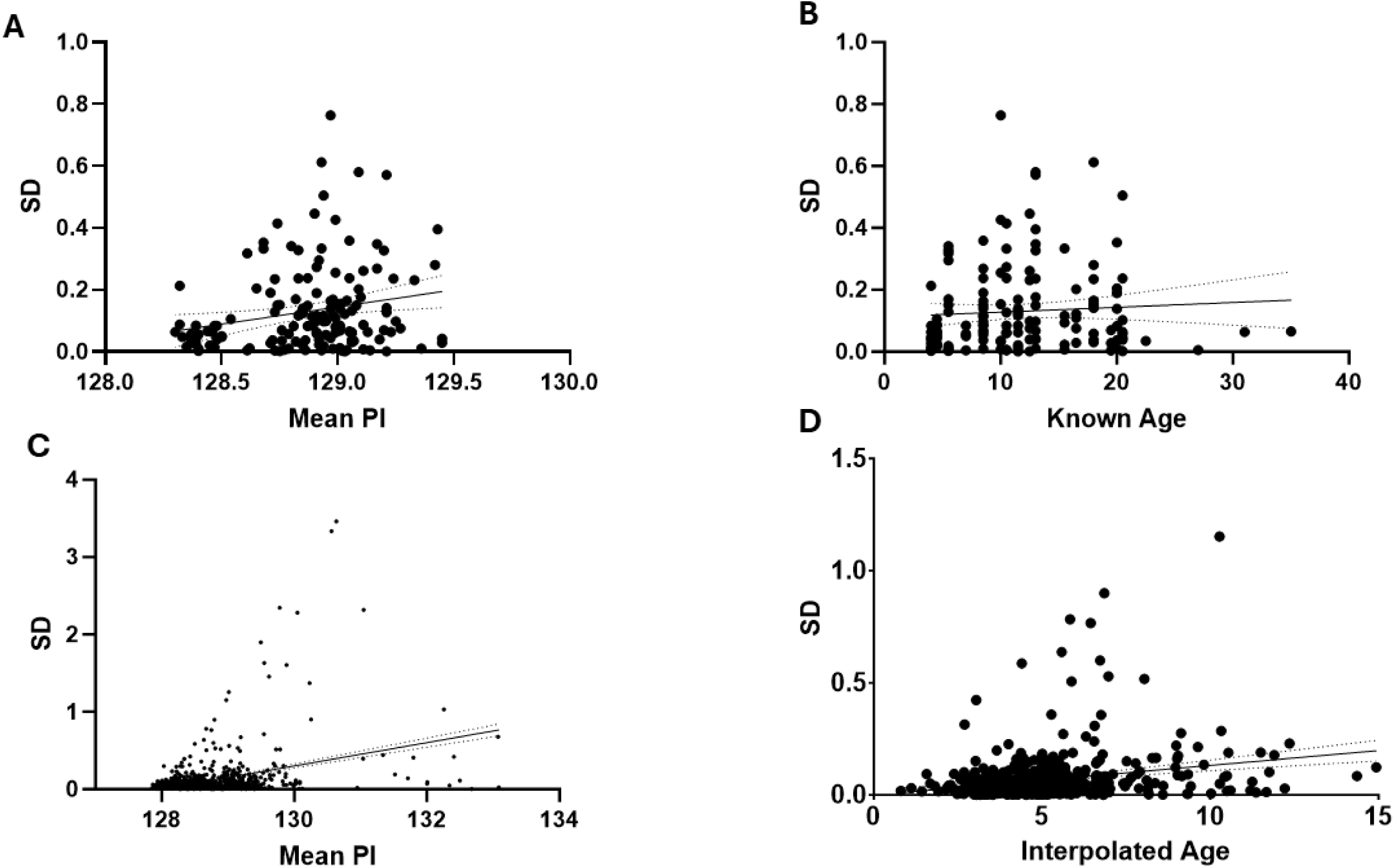
The standard deviation of the mean pixel intensity values for each set of a mosquito’s wings is modestly correlated with increasing mean pixel intensity (PI) or mosquito age. Each dot represents data from a set of a mosquito’s wings. Top panels: laboratory-reared *An. gambiae* aged in a mesocosm (n=149 in both A and B). Bottom panels: *An. gambiae* s.l. captured during the RIMDAMAL II trial (C; n =1861), (D; n = 549). Regression line slopes in panels A, C, and D are significantly different from zero (panel A, P=0.008, r2 = 0.045; panel C, P< 0.0001, r2 = 01474; panel D P< 0.0001, r2 = 0.054)

### Individual and binned pixel intensity of mosquito populations to examine inferred age structure

Next, we plotted the proportions of wild mosquitoes from the RIMDAMAL II trial from which we took wing pictures, both individually and in bins of 0.3 PI units, to observe the overall inferred age structure (Figure 2). The plots shows distributions that mimic wild mosquitoes age-graded by the gold standard Polovodova method ^5^ as well as newer techniques such as MIRS ^23^, suggesting a population distribution dominated by nulliparous and young mosquitoes in the lowest PI ranges, and relatively few older females in the long top-or right-skewed tail of the individual or bin distributions, respectively (Figure 2). Overall, most mosquitoes captured during the trial had PIs of ≤ 128.73 (Range: 127.81-133.43) with a top-or right-skewed distribution tail extending to a PI of 133.43. When the population was split based on mosquitoes captured in each arm of the trial, the top-or right-skewed tail of the distribution was more robust among the population from placebo clusters but not intervention (ivermectin-treated) clusters (Figure 3). Furthermore, when separating the data into the trial years, the top-or right-skewed tail of the distribution was evident among the population captured in 2019 but absent in 2020 (Figure 4). The PI bin distributions among the populations captured in 2020 among clusters from both arms were grouped in only 3 of the lowest PI bins (Figure 4). A generalized linear mixed model was generated using individual wing PIs from these mosquitoes, using a gamma distribution with a log link function and a random (grouping) mosquito identification term. This model revealed a small but significant reduction in PI of mosquitoes captured in 2020 compared to 2019 (Ratio (exponentiated model coefficient): 0.9955 [95% CI 0.9950 – 0.9961], p < 0.001). No significant difference was detected between control and treatment arms in either 2019 (p = 0.31) or 2020 (p = 0.64).

**Figure 2.**
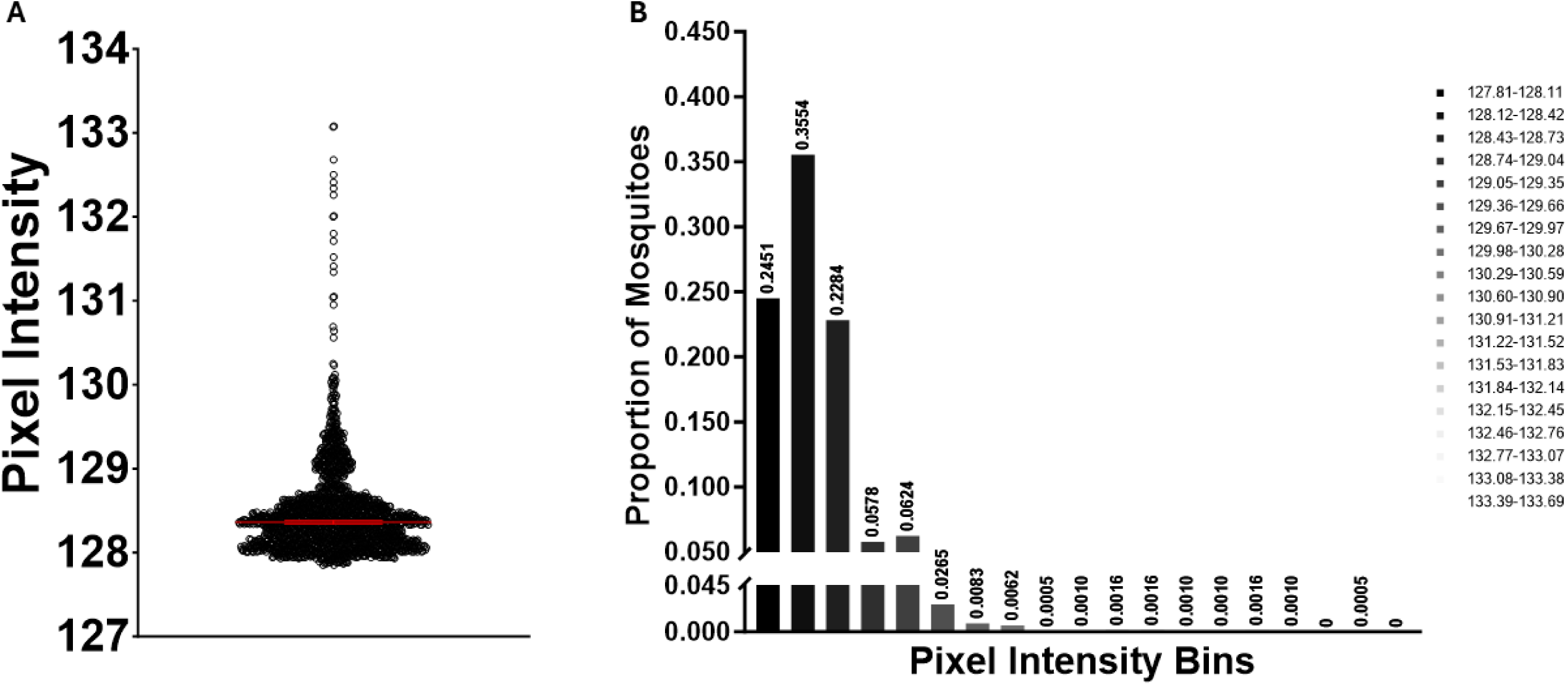
Pixel intensities (PI) from wild *Anopheles gambiae* s.l. captured in the RIMDAMAL II trial reflects typical population age distributions from wild mosquitoes. A) The distribution of individual mosquito’s mean wing PIs. Each dot represents the mean PI from a set of a mosquito’s wings. B) Bins of mosquitoes’ mean wing PIs by 0.3 PI units/bin.

**Figure 3.**
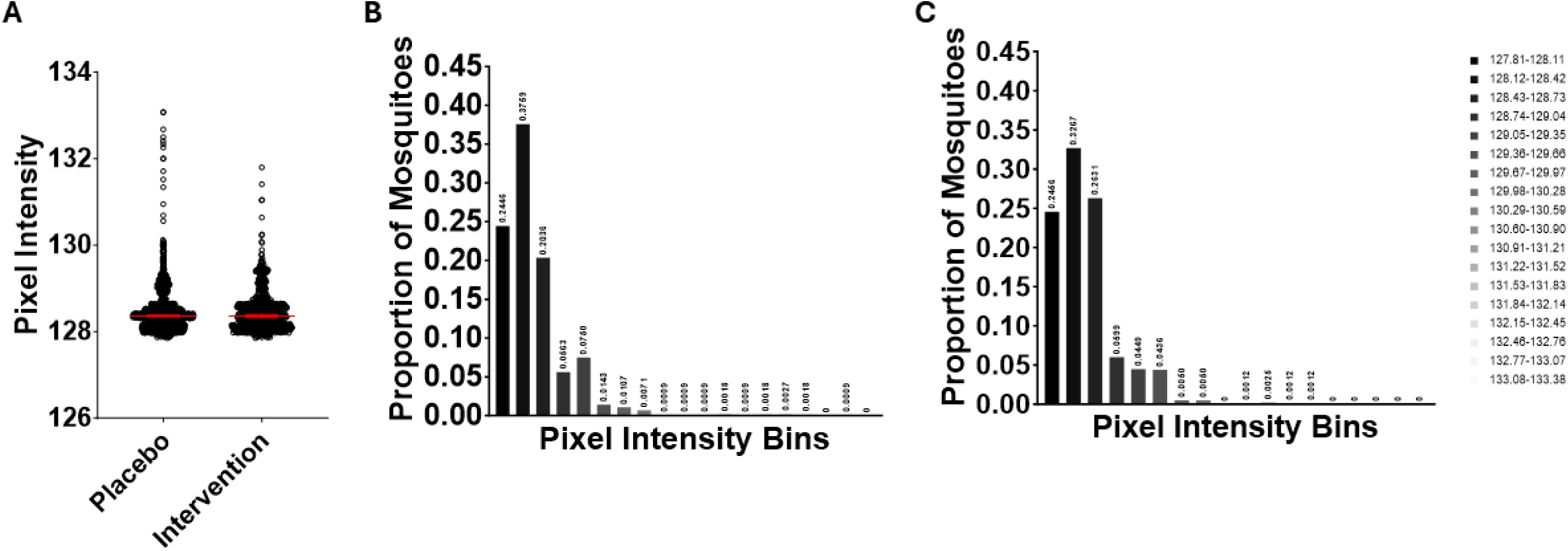
Pixel intensities (PI) of wild *Anopheles gambiae* s.l. captured in the RIMDAMAL II trial, grouped by arm. A) The distribution of individual mosquitoes’ mean wing PIs from the Placebo and Intervention arms. Each dot represents the mean PI from a set of a mosquito’s wings. B) Bins of mosquitoes’ mean wing PIs, from mosquitoes captured from clusters of the Placebo (B) or Intervention (C) arms.

**Figure 4.**
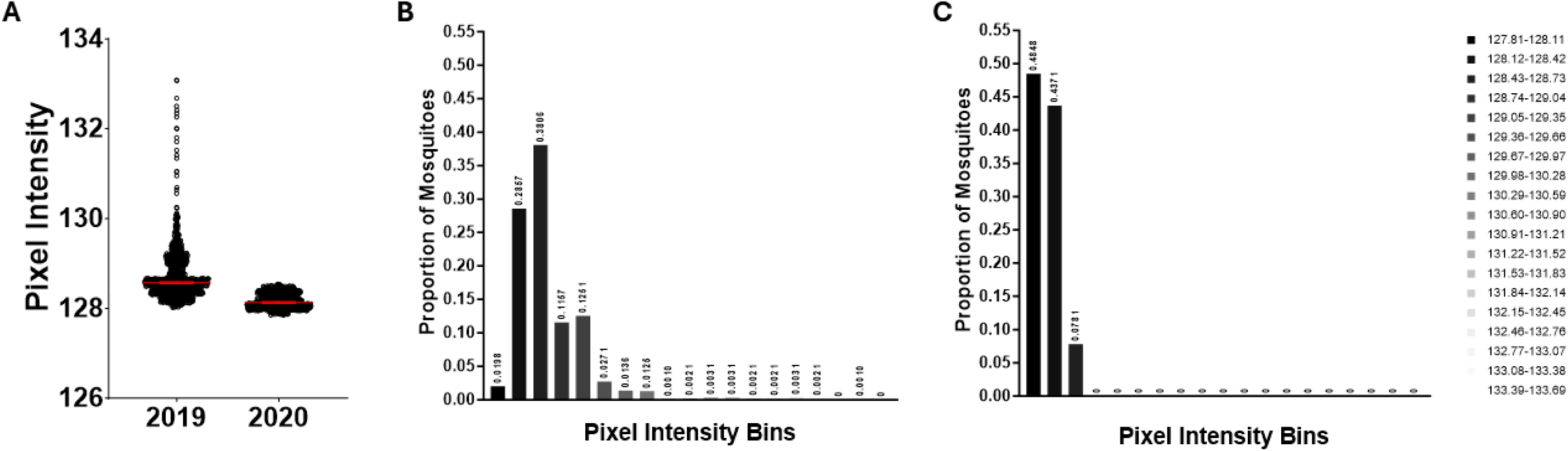
Pixel intensities (PI) of wild *Anopheles gambiae* s.l. captured in the RIMDAMAL II trial, grouped by trial year. A) The distribution of individual mosquitoes’ mean wing PIs grouped by the year they were captured during the rainy season intervention periods (late July to early November of 2019 and 2020). Each dot represents the mean PI from a set of a mosquito’s wings. B) Bins of mosquitoes’ mean wing PIs, from mosquitoes captured during trial years 2019 (B) and 2020 (C).

These PI distributions associate with the timings of the two mosquito control interventions that occurred during the trial, whereby ivermectin or placebo was distributed in the respective trial arms throughout both years of the trial, and distribution of new dual chemistry Interceptor® G2 (IG2) nets (containing chlorfenapyr + alpha-cypermethrin) occurred in clusters of both arms in early November 2019, towards the very end of the first rainy season approximately 2-4 weeks after the 4^th^ MDA between sampling weeks 13 and 15 when mosquito capture numbers were very low (Figure 5)^28^. Thus, most of the IG2 nets were not deployed by the populace until 2020 and were used by households from both arms throughout that year. This association can be observed when comparing the median PIs per mosquito population collected one and three weeks after each MDA performed in the trial (Figure 5). In 2019, the median PI among *An.* gambiae s.l. caught in placebo clusters three weeks after each of the first two MDAs were significantly higher relative to the populations captured one week after those MDAs (Sampling period 1 vs 2, p ≤ 0.01; sampling period 3 vs 4, p ≤ 0.001), while the median PIs significantly decreased from populations caught in ivermectin-treated clusters over the same sampling time frames (Sampling period 1 vs 2, p ≤ 0.01), suggesting an ivermectin effect on mosquito population PI structure that manifests by the third week after the 1^st^ and 2^nd^ MDAs (Figure 6). This ivermectin effect on wing PI then became inapparent after the 3^rd^ MDA in 2019 and through all MDAs in 2020 when the IG2 nets were used throughout the 2020 season. Overall, the new nets seemed to have a dramatic effect on the median PI, lowering it below 128.5 among mosquito populations of both arms beginning with the 4^th^ MDA and lasting to the end of the trial.

**Figure 5:**
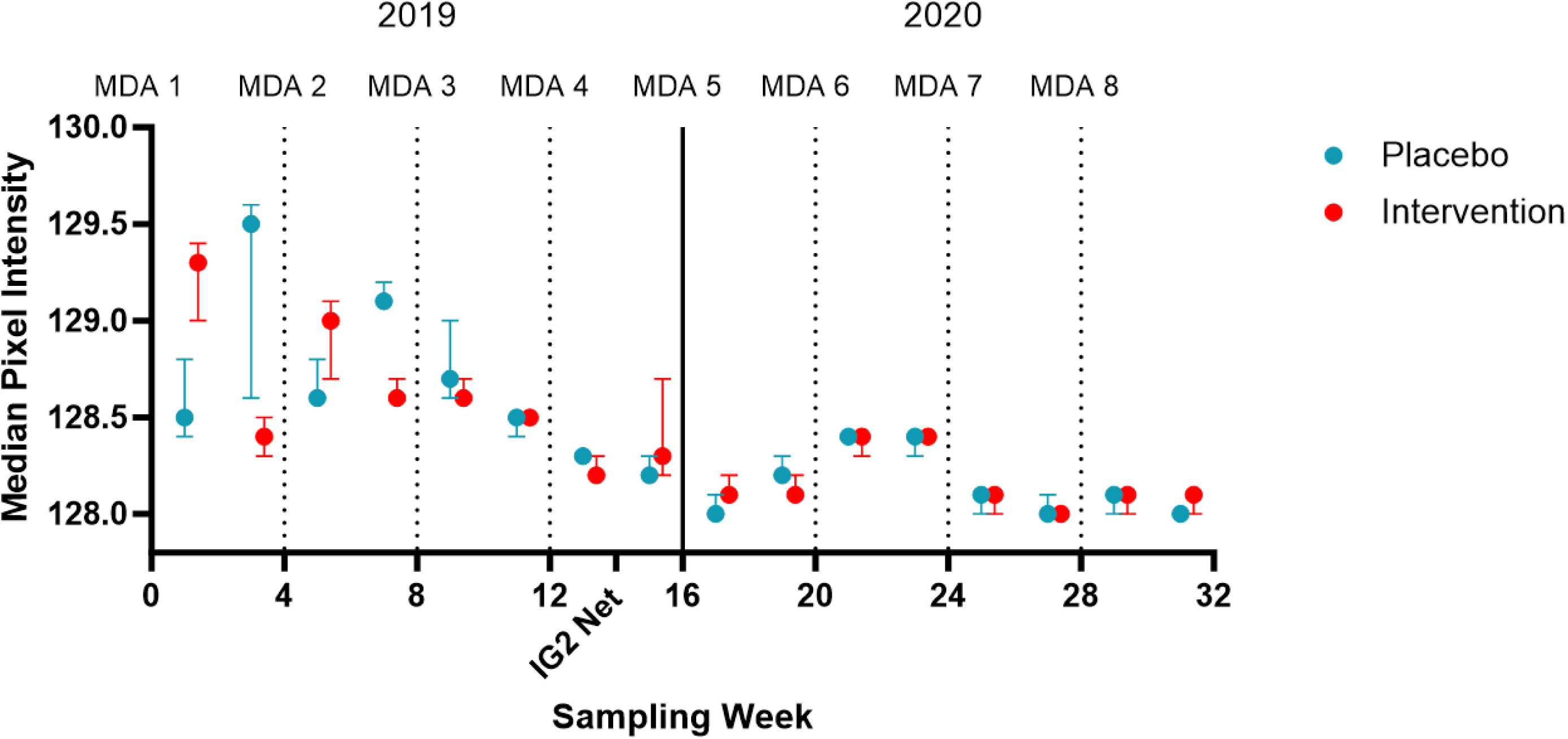
Median pixel intensities (PIs) from wild *Anopheles gambiae* s.l. captured in the RIMDAMAL II trial by sampling period and relative to the timing of the mass drug administrations. Median PI of mosquitoes captured from the Placebo (blue: n=1129) and Intervention (red: n=799) arms. Whiskers depict 95% CIs. Mosquito sampling weeks over the 2-year intervention period (late July-early November in 2019 and 2020) are enumerated consecutively on the X-axis, and mass drug administrations (MDA) of either placebo or ivermectin tables are marked with vertical or dotted lines and occurred over a 3-day periods. The period of IG2 net distributions in 2019 to all villages in the health district is marked on the X-axis.

**Figure 6:**
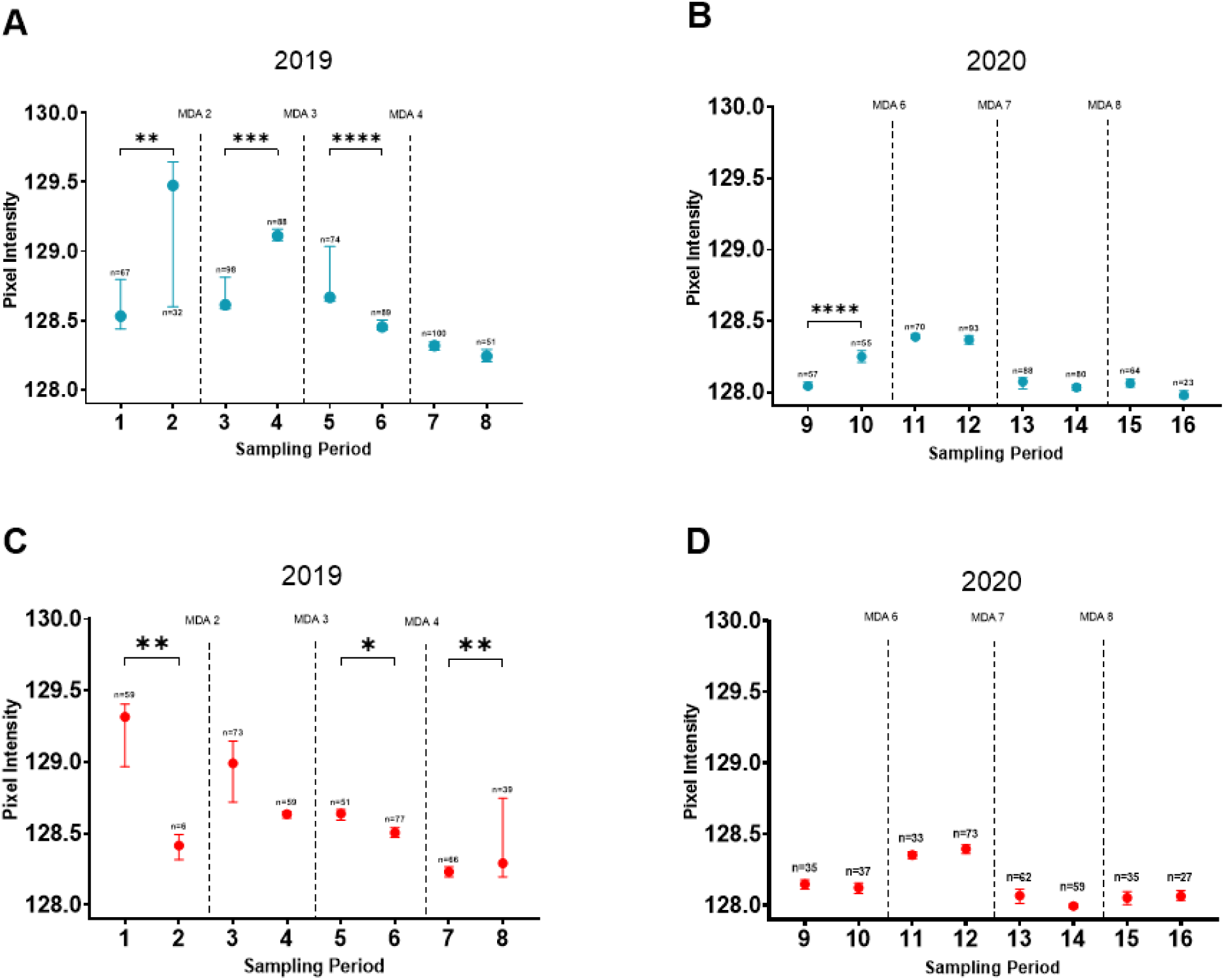
Comparisons of median pixel intensities (PIs) from wild *Anopheles gambiae* s.l. captured in the RIMDAMAL II trial by sampling period and relative to the timing of the mass drug administrations. Median PI of mosquitoes captured from each arm were compared by between the two populations sampled 1-week and 3-weeks following each mass drug administration (MDA). Data from mosquitoes captured in the Placebo (blue: n=1129) and Intervention (red: n=799) arms are showed in panel A & B and C & D, respectively. Whiskers depict 95% CIs. Mosquito sampling weeks over the 2-year intervention period (late July-early November in 2019 and 2020) are enumerated consecutively on the X-axis, and mass drug administrations (MDA) of either placebo or ivermectin tables are marked with vertical or dotted lines and occurred over a 3-day periods. Comparisons were ANOVAs and post hoc Kruskal-Wallis tests; (*= P ≤ 0.05; **= P ≤ 0.01; ***= P ≤ 0.001; ****= P ≤ 0.0001).

### Adapting and applying an age model to wild An. gambiae s.l

To interpolate age for the wild mosquitoes, we sought to use the colonized mosquitoes aged in laboratory mesocosms to develop an age model and use it to predict the ages of the wild mosquito populations from RIMDAMAL II^27^. The range of mean PI scores from these lab mosquitoes of known age was notably constrained (range: 128.30 – 129.45) compared to the range obtained from wild mosquitoes (range: 127.85 – 133.08), but the agreement in the magnitude of PI measured in two the different settings and between the lab and field populations was encouraging. The lowest mean PI score from the newly eclosed adults (128.35) was 0.5 PI units higher than the lowest mean PI score obtained from wild mosquitoes (127.85), suggesting wings from the wild mosquitoes tended to have more and/or darker scales than those from the lab colony, or that wing pictures from the microscope camera in the field were more variable than those in the lab (perhaps due to different lighting conditions, etc.).

A simple linear regression model was first applied to the lab mosquitoes, which had relatively poor fit to the data (*r*^2^ = 0.4011) and predicted the wild mosquitoes to be between –24.31 and 163.17 days old. We then performed stepwise comparisons of multiple non-linear models that may plausibly fit the data and reflect the biology of scale loss over time. The best fit model was a symmetrical sigmoidal curve with a variable slope (difference in AICc from the linear model = 90.33; *r*^2^ = 0.6577), from which the best-fit values were PIs between 128.4 (95% CI 128.3 – 128.5) and 129.0 (95%CI 129.0 – 129.1). With this model, PIs from 128.30-128.55 were predicted for mosquitoes between <1-4 days old, PIs from 128.56-128.96 were predicted for mosquitoes 5-9 days old, and PIs from 128.97-129.45 were predicted for mosquitoes ≥10 days old (Figure 7). However, given the variance of wild mosquito PI data compared to the model, predicted age could only be accurately interpolated from approximately 30% (578/1928) of the wild mosquitoes (Figure 8). Nonetheless, using this model, we binned the mosquitoes into the 3 interpolated pixel intensity groups and observed a slight but non-significant difference in the structure of these binned mosquitoes in 2019, whereby those in from the placebo arm had fewer ‘middle aged’ mosquitoes with PIs between 128.57-128.96 and more ‘old’ mosquitoes with PIs between >128.96 in the ivermectin arm. However, the difference in PI bin structures in 2020 compared to 2019 were noticeable, whereby all mosquitoes tested were predicted to be ‘young’ (PI < 128.57) in both arms (Figure 9).

**Figure 7.**
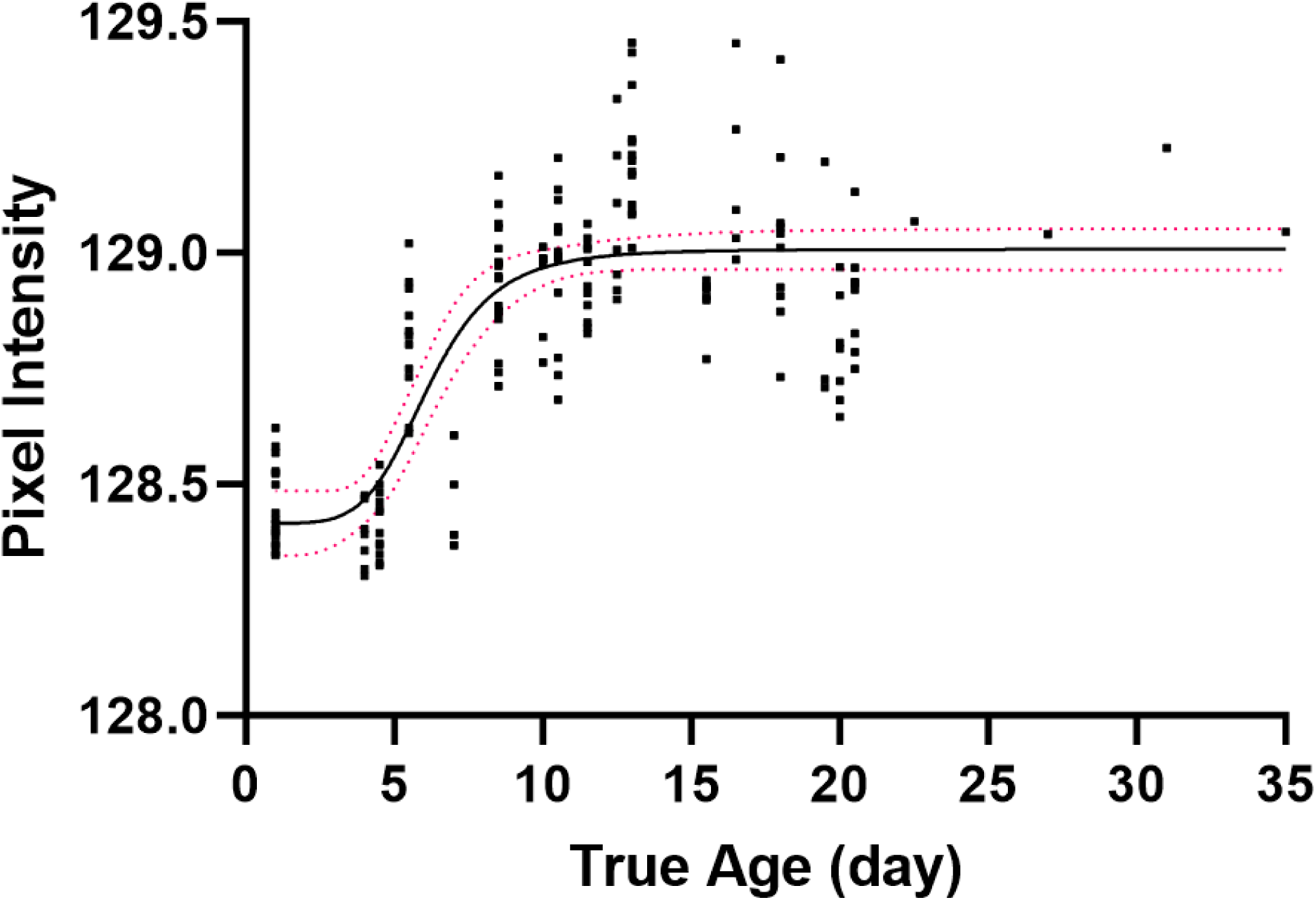
Age model derived from colonized, marked *Anopheles gambiae* reared in laboratory mesocosms to known ages. Mean pixel intensities of colonized mosquitoes reared to known ages (n=169) were plotted by their true age and fitted by stepwise comparisons to a linear and multiple non-linear models. The best fit model (depicted) was a variable slope sigmoidal model. The best fit line of the final model is in black and red dotted lines represent 95% confidence bands.

**Figure 8.**
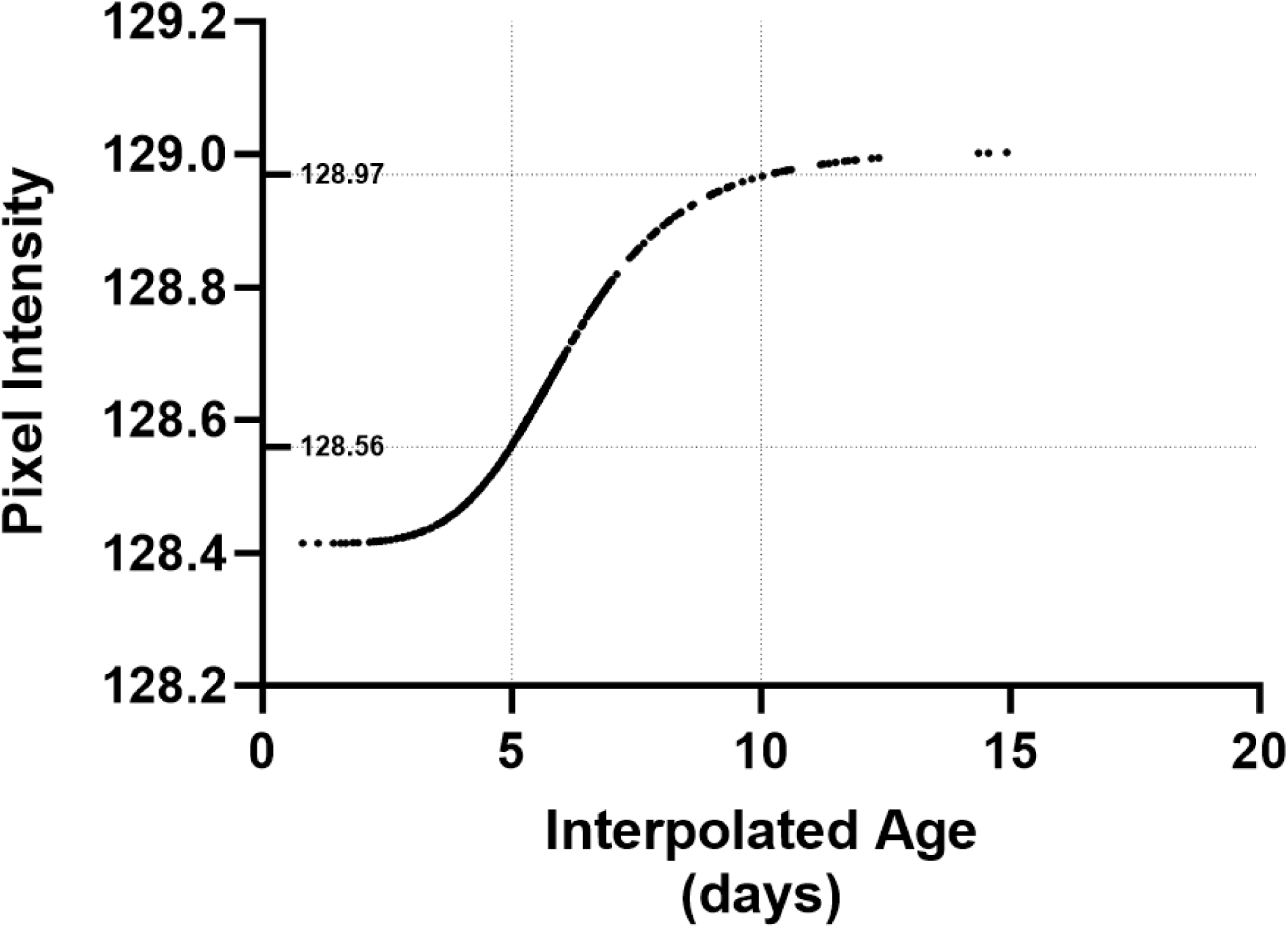
Interpolated ages of wild *Anopheles gambiae* s.l. captured in the RIMDAMAL II trial using the pixel intensity-based age model derived from colony mosquitoes. Dots are individual wild mosquitoes placed on the best-fit-line of age model; 30% (n=578) of the wild mosquitoes’ pixel intensity (PI) values fit within the model’s lowest and highest range of 95% CIs. PIs corresponding to the interpolated ages of 5 and 10 were plotted (dotted lines) using the model.

**Figure 9.**
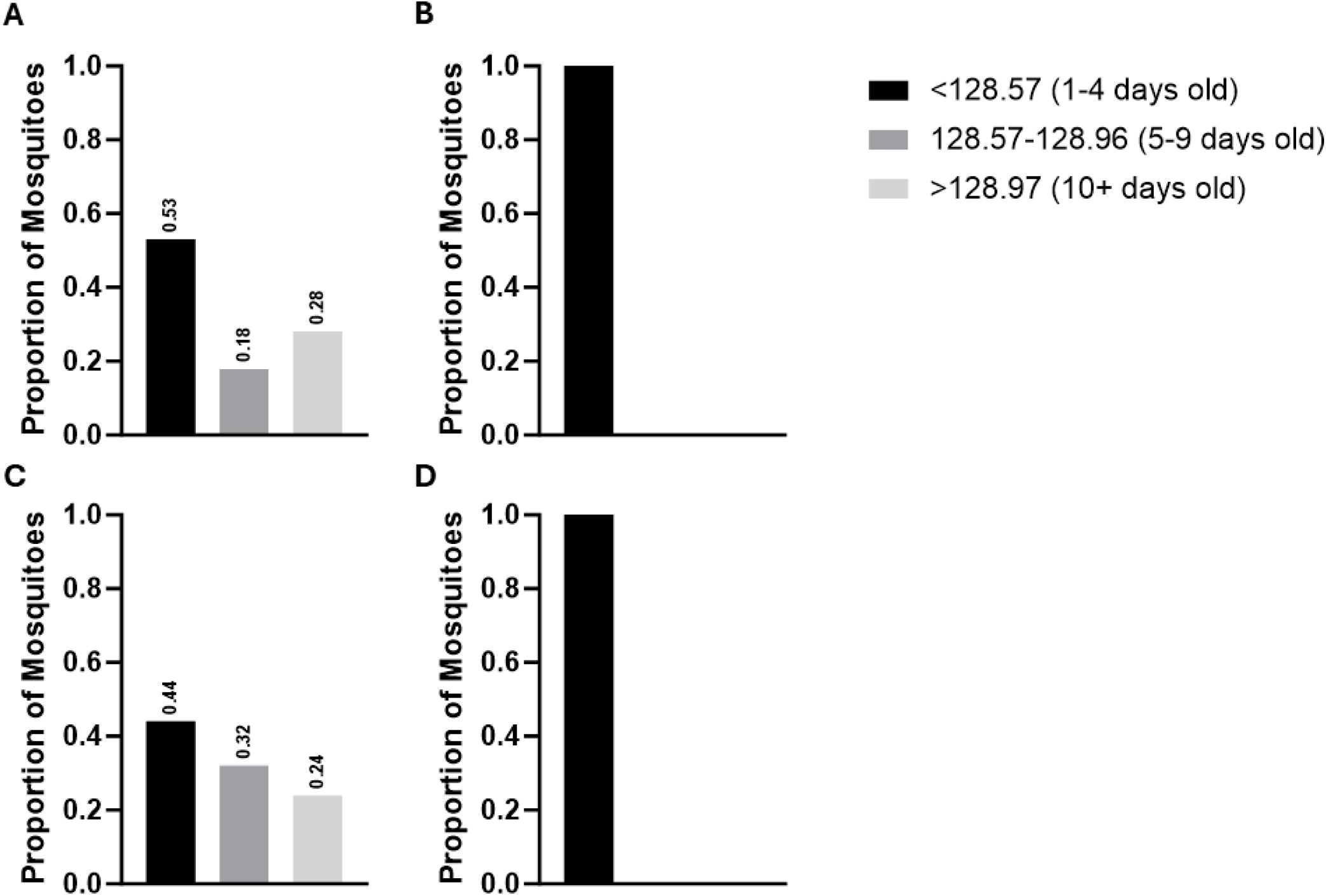
Wild *Anopheles gambiae* s.l. captured in the RIMDAMAL II trial grouped in into three, pixel intensity (PI) bins corresponding to interpolated age classes. A) mosquitoes from the first year of the trial (2019) and caught in Placebo clusters. B) mosquitoes from the second year of the trial (2020) and caught in Placebo clusters. A) mosquitoes from the first year of the trial (2019) and caught in Intervention clusters. A) mosquitoes from the second year of the trial (2020) and caught in Intervention clusters.

## Discussion

Here we demonstrate that this revised PI analysis of wing photos enables normalization of the data collected and that the PI distributions, either from individual mosquitoes or with binned data, are similar to mosquito age structures observed from using the gold standard Polovodova method consisting of ovarian dilation counting, which is sometimes combined with analysis of Christophers stages^11,23,29–32^.

Furthermore, we show how this method can be used to observe changes in mosquito population structure in response to two different vector control methods that were performed during the RIMDAMAL II trial, specifically ivermectin mass drug administrations and use of IG2 nets.

The most prominent changes in PI distributions from *An. gambiae* s.l. captured at intervals during the RIMDAMAL II trial were observed in the differences between intervention seasons in 2019 and 2020. These changes were associated with the distribution of new IG2 nets at the end of the 2019 season, just prior to the last mosquito sampling period of that year when the rains had ended and the populations of *An. gambiae* s.l. were nearly gone from the study site. The efficacy of IG2 nets on wild *An. gambiae* has been thoroughly examined, and is known to strongly effect mosquito populations^33–36^. The entomological indices we previously reported from the trial suggested a similarly strong effect on malaria vectors from both arms of the trial and that occurred primarily in the in trial’s the second season (2020)^28,37^. This effect included, a) significant mortality (∼50%) of indoor-captured, blood fed mosquitoes held in survival bioassays from both arms, and b) significant reductions in the number of the *An. gambiae* s.l captured in 2020, which were approximately half of those captured in 2019, and a near absence of *An. funestus* captured at the study site in 2020 (n = 49) relative to 2019 (n = 1148) despite similar rainfall and climate at the site across these two years. Similarly, the PI analysis reported from mosquitoes captured in 2020 relative to 2019 (Figure 9) supports the strong mosquito killing effect from these nets and demonstrates how PI can be used to track the efficacy of a robust malaria vector control method like IG2 nets.

The ability of this age-grading method to detect mosquito population changes from the effect of ivermectin mass drug administrations is more nuanced, but the ivermectin intervention itself was not successful in reducing malaria incidence in the trial, nor in reducing the classical entomological indices of density and entomological inoculation rate^28^. The primary entomological index that was affected in treatment clusters compared to placebo clusters was the wild blood fed *An. gambiae* s.l. survival rate, but only in 2019 and only from blood fed mosquitoes captured in the week following the MDAs. Regarding the PI of these mosquito populations, there were modest detectable differences between treatment and control arms at the beginning of the trial in 2019 observed from the mosquito sampling periods that occurred three weeks after MDAs 1-3 were administered. It is interesting that these PI trends mirror the trends of infectious *An. gambiae* s.l. bites per person per night in each arm shown over the trial, suggesting that lower median PIs may correlate with fewer infectious mosquitoes in the population^28^. The observation that the strongest effect of ivermectin MDA on mosquito PI seems to be observed 3 weeks after the MDAs occurred, when the direct mosquitocidal effects are inapparent and the drug is at very low pharmacokinetic levels or undetectable in most of the blood samples tested from the populace, suggests a delayed action of the drug’s effect on mosquito population structure and infectiousness^28,38^. Perhaps this delayed effect is due to increased probability of killing the fewer, older mosquitoes that were actively blood feeding immediately after the MDA was administered, and leaving unharmed most of the younger, newly emerged mosquitoes with lower PIs and that have yet to blood feed.

The age model developed using PIs obtained for mesocosm-aged laboratory colony of *An. gambiae* was somewhat limited due to potential differences in the mosquitoes’ wings, as well as the laboratory vs. field conditions experienced by the mosquitoes and possibly also the wing imaging performed. Overall, the lab mosquitoes had a very narrow PI range that limited our ability to interpolate age from wild mosquitoes with PIs residing outside the range of our model. Nonetheless, the best fit symmetrical sigmoidal model that fit the PI data from these laboratory mosquitoes and was used predict the age of the wild mosquitoes from the trial suggests little differences in the PIs among very young (1-4 days old) and very old (≥10 day old) mosquitoes, respectively, but a rapid shift from between these young to old classes in between the ages of 4 to 10 days, presumably due to extensive flying and loss of wing scales during this timeframe. Importantly, wild-type *Anopheles gambiae* do not become infectious until they complete the minimum EIP, which typically correlates with them reaching the 2-or 3-parous state, or approximately 8-11 days old, thus our model’s ability to predict proportions of ‘old’ mosquitoes with cut-off PIs ≥129.7 may be a simple method to classify potentially infectious *An. gambiae* in a population^4,5,30^. Nonetheless, it remains to be seen if this model will similarly fit PI data from wild type mosquitoes reared in large outdoors mesocosms in Africa.

While our newer analysis method allowed for better normalization of wing photos, further work could be done to refine the use of wing pictures to age mosquitoes. Technologies which auto-crop images to only include the wing, or which focus PI or machine vision analyses on specific wing areas that are more sensitive to scale loss from flying (such as the wing fringe), could lead to a more refined analysis of mosquito age structures. Additionally, further standardization of photo backgrounds and microscope light intensities would likely add more control to the method and allow for easier comparisons across space and time. We believe that these refinements would abrogate the need for regionally calibrated age models at each place and time such analyses were to occur. Finally, due to sample preservation and costs, we were not able to speciate most of the mosquitoes from the entire *Anopheles gambiae* sensu lato complex from RIMDAMAL II. As such, there may be differences in *An. gambiae* sensu strictu, *An. coluzzi* and *An. arabiensis* PIs^28,37^. Therefore, further investigations into the *An. gambiae* species complex are needed to determine if specific species have differing PI ranges.

Overall, we have demonstrated how using PI on wing images can be used to easily age grade *Anopheles gambiae* mosquito populations. This method might easily be expanded to allow investigators and vector control experts across the world, who are implementing and evaluating control of malaria, helminth and arbovirus vectors, to expediently determine mosquito population ages from their surveillance efforts. Such efforts may be highly valuable for evaluation of their control applications and estimations of the risk of mosquito-borne disease spread in their communities.

## M&M

### Mosquito rearing

*Anopheles gambiae* G3 were reared at 27-30°C with 60-80% relative humidity. A standard photoperiod of 16 hours light:8 hours dark was utilized. Larvae were raised in open-top plastic bins and fed a diet of TetraMin fish food (Spectum Brands Pet, LLC). Pupae were transferred into closed-top plastic bins. 24 hours after emergence, adults were aspirated and placed into a closed-top plastic adult bin. Adults were grouped based on their respective date of emergence to ensure adults were the same age. If adult collection was limited, groups were mixed but never exceeded more than two days in age difference.

### Mesocosm experiment

These experiments were well described in Gray *et al*, 2022. In brief, 200 *An. gambiae* G3 mosquitoes were marked using florescent pigment powder (Shannon Luminous Materials, Inc.) placed into a large (122 cm long x 61 cm wide x 96.5 cm high) plastic bin in which a fake plant was placed in the middle to force the mosquitoes navigate around as they sought water and bloodmeals that were placed on opposite sides of the bin. Plastic dishes with holes cut into them covered the oviposition papers and sugar cubes. These obstacles were used to replicate a ‘wild’ environment for the lab-reared mosquitoes to force flying behaviors. Twice a week, 15 mosquitoes were removed from the mesocosm, and their wings were detached for analysis.

### RIMDAMAL II mosquito collection and photo acquisition

During the RIMDAMAL II clinical trial, entomological sampling was conducted to compare entomological indices, such as bioassay survival, entomological inoculation rate, and age structure, between the control and treatment arms of the trial. Mosquitoes were collected from 6 (3 treatment; 3 placebo) clusters (villages or village sectors), 1 and 3 weeks after each mass drug administration using a Prokopac aspirator inside predetermined households. Mosquitoes were then transported to the field station, speciated, and *Anopheles* mosquitoes were retained. A portion of *An. gambiae* s.l. mosquitoes were subjected to wing removal. Removed wings were placed on a glass slide and photographed using a Leica EZ4 stereoscope (Leica Microsystems, USA). Photos were then electronically transmitted to Colorado State University for pixel intensity (PI) analyses.

### Digital wing photo analysis

Each mosquito had photos taken of both wings using same Leica EZ4 stereoscope and PI was calculated for each wing using the wing photos processed by the BigPicture program developed by Viden Technologies, LLC. Subsequently, the mean of the left– and right-wing PIs were calculated to establish a PI per individual mosquito (a single PI from the mean of both wings). If a mosquito had only one wing image, or if one wing was folder or torn during the slide mounting process, the individual mosquito was given a PI that came from only one of its wings.

### Statistical and mathematical methods

Raw PI values from the trial were modeled using a log-linked gamma generalized linear model. Left– and right-wing PI values were input into the model individually, with a mosquito-level random effect term as a grouping variable. Fixed effects of treatment arm, year, and their interaction term were included. To delineate population PI range differences, mosquitoes were placed into bins with a difference of 0.3 PI. Furthermore, mosquitoes were separated based on the arm location they were captured in to determine differences in PI bins. To establish an age-to-pixel intensity model, mean PIs from mesocosm-reared colonized mosquitoes were first analyzed with a linear model using GraphPad Prism (v10.1.1), and then in a stepwise approach all non-linear models in the program were compared to the linear model using Akaike’s Information Criterion. The best fit model emerging from this process was Sigmoidal Variable Slope model. The predicted ages of the wild mosquitoes captured in the RIMDAMAL II trial were then interpolated using this best-fit model.

## Figures and Tables

**Table 1:**
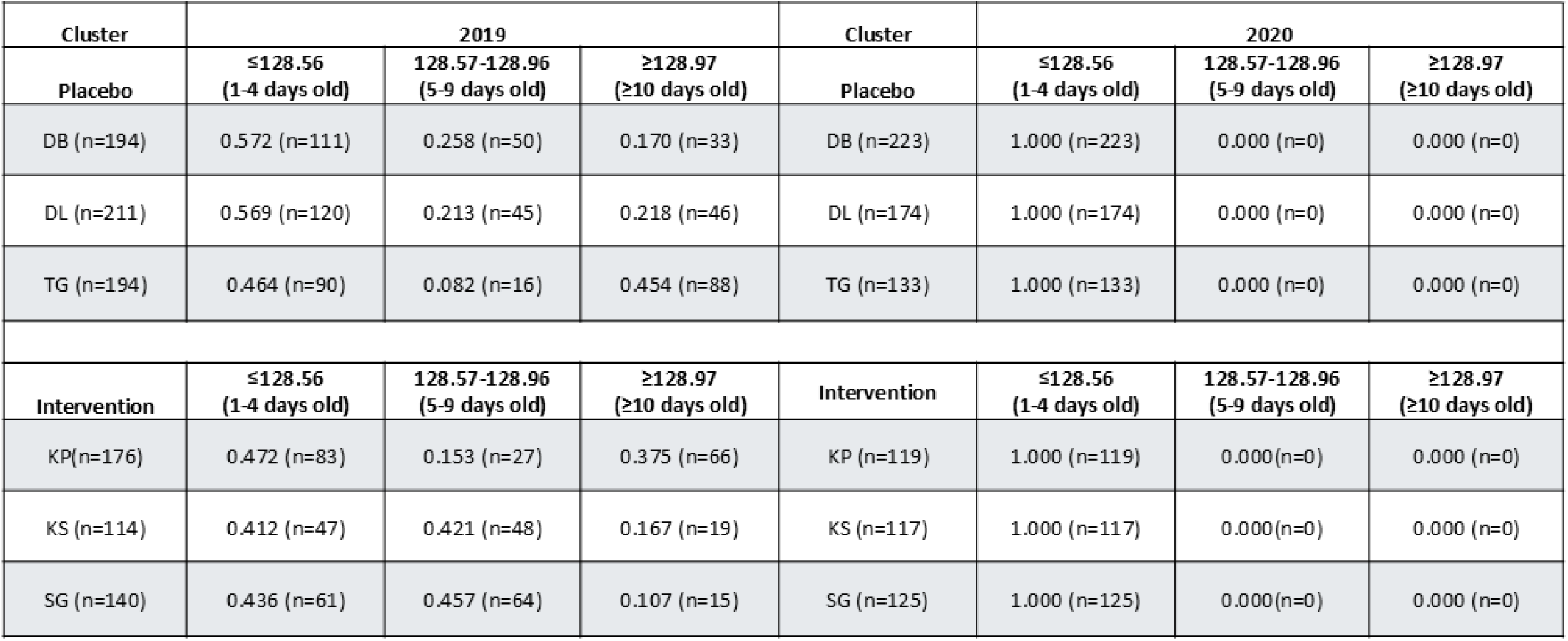
Proportions and totals number of wild *Anopheles gambiae s.l*. captured from each RIMDAMAL II cluster and arm, and during each year, corresponding to their pixel intensities and associated with their interpolated age class in days.

**Supplemental Table 1.** Times of mass drug administrations and mosquito sampling periods and weeks associated with the RI**MDAMAL** II trial.

|  |  |  |  |  |
| --- | --- | --- | --- | --- |
| **MDA** | **Sample Period** | **Sample Week** | **Date Range** | **Year** |
| MDA 1+1 | 1 | 1 | July 30 - Aug 3 | 2019 |
| MDA 1+3 | 2 | 3 | Aug 13 - 15 | 2019 |
| MDA 2+1 | 3 | 5 | Aug 27 - 29 | 2019 |
| MDA 2+3 | 4 | 7 | Sept 11 - 13 | 2019 |
| MDA 3+1 | 5 | 9 | Sept 25 - 27 | 2019 |
| MDA 3+3 | 6 | 11 | Oct 9 - 12 | 2019 |
| MDA 4+1 | 7 | 13 | Oct 21 - 26 | 2019 |
| MDA 4+3 | 8 | 15 | Nov 7 - 11 | 2019 |
| MDA 5+1 | 9 | 17 | Jul 29 - Aug 3 | 2020 |
| MDA 5+3 | 10 | 19 | Aug 12 - 17 | 2020 |
| MDA 6+1 | 11 | 21 | Aug 28 | 2020 |
| MDA 6+3 | 12 | 23 | Sept 7 - 13 | 2020 |
| MDA7+1 | 13 | 25 | Sept 21 - 23 | 2020 |
| MDA7+3 | 14 | 27 | Oct 7 - 12 | 2020 |
| MDA 8+1 | 15 | 29 | Oct 20 - 22 | 2020 |
| MDA 8+3 | 16 | 31 | Nov 6 | 2020 |

**Supplemental Figure 1.**
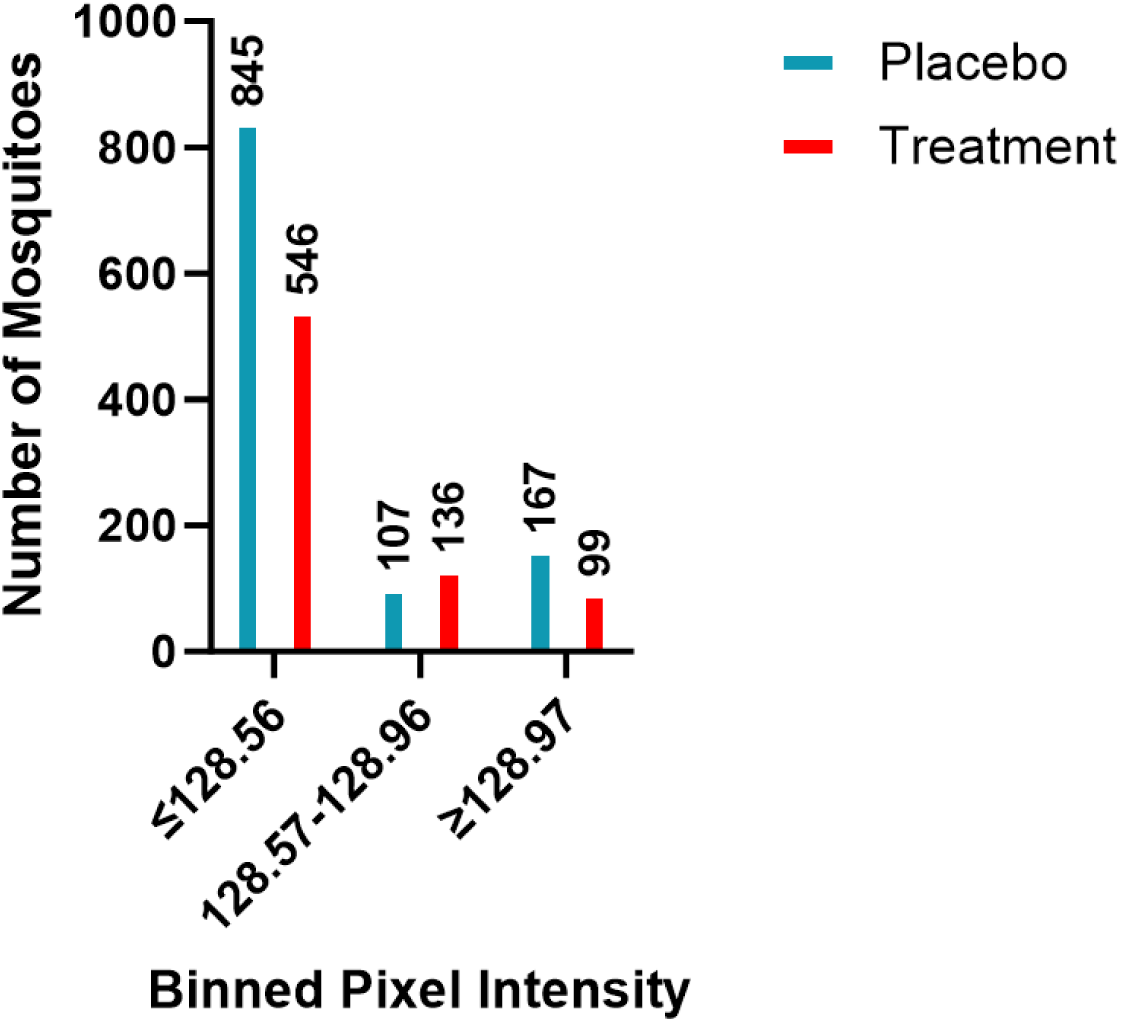
Numbers of wild *Anopheles gambiae* s.l. captured in the RIMDAMAL II trial and grouped by pixel intensity and trial arm.

**Supplemental Figure 2.**
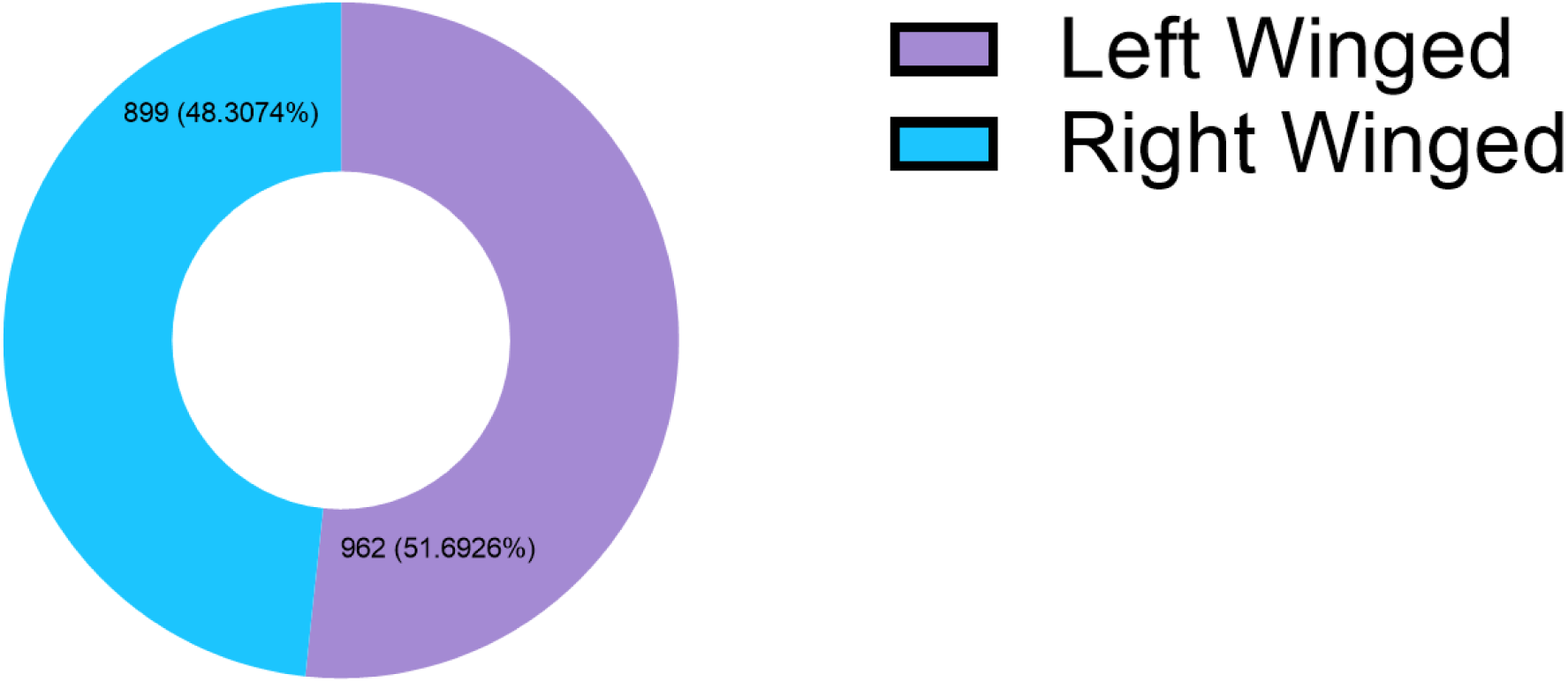
Mosquito populations do not have a favored wing. The difference between the left and right pixel intensities were calculated. A positive difference was determined to be favorable to the left wing which a negative difference was favorable to the right wing. A binomial test found there was no significance between the two populations with the assumption that wing favorability would be evenly split within the population indicating the population does not have a favored wing.

## References

1. Charlwood, J. D., Pinto, J., Sousa, C. A., Ferreira, C. & Petrarca, V. ‘A mate or a meal’ – Pre-gravid behaviour of female Anopheles gambiae from the islands of São Tomé and Príncipe, West Africa. Malaria Journal 11 (2003).

2. Ohm, J. R. et al. Rethinking the extrinsic incubation period of malaria parasites. Parasites Vectors 11, 178 (2018).

3. Guissou, E. et al. A non-destructive sugar-feeding assay for parasite detection and estimating the extrinsic incubation period of Plasmodium falciparum in individual mosquito vectors. Sci Rep 11, 9344 (2021).

4. Gillies, M. T. & Wilkes, T. J. A study of the age-composition of populations of Anopheles gambiae Giles and A. funestus Giles in North-Eastern Tanzania. Bulletin of Entomological Research 56, 237–262 (1965).

5. Ryan, S. J., Ben-Horin, T. & Johnson, L. R. Malaria control and senescence: the importance of accounting for the pace and shape of aging in wild mosquitoes. Ecosphere 6, art170 (2015).

6. Perry, M. Malaria in the Jeypore Hill tract and adjoining coastland. Paludism 5 32–40 (1912).

7. Gillett, J. D. Age Analysis in the Biting-Cycle of the Mosquito Taeniorhynchus (Mansonioides) Africanus Theobald, based on the Presence of Parasitic Mites. Annals of Tropical Medicine & Parasitology 51, 151–158 (1957).

8. Detinova, T. S. AGE-GROUPING METHODS IN DIPTERA OF MEDICAL IMPORTANCE. 213 (1962).

9. Detinova, T. S. & Gillies, M. T. Observations on the Determination of the Age Composition and Epidemiological Importance of Populations of Anopheles gambiae Giles and Anopheles funestus Giles in Tanganyika.

10. Polovodova, V. P. Changes with age in the female genitalia of Anopheles and the age composition of mosquito populations. Moskva (Thesis) (1947).

11. Beklemishev, W. N., Detinova, T. S. & Polovodova, V. P. Determination of Physiological Age in Anophelines and of Age Distribution in Anopheline Populations in the USSR. Bull. Wld Hlth Org. 21, 223–232 (1959).

12. Lambert, B. et al. Monitoring the Age of Mosquito Populations Using Near-Infrared Spectroscopy. Sci Rep 8, 5274 (2018).

13. Cook, P. E. et al. The use of transcriptional profiles to predict adult mosquito age under field conditions. Proc. Natl. Acad. Sci. U.S.A. 103, 18060–18065 (2006).

14. Sikulu, M., et al. Near-infrared spectroscopy as a complementary age grading and species identification tool for African malaria vectors. (2010).

15. Hugo, L. E. et al. Proteomic Biomarkers for Ageing the Mosquito Aedes aegypti to Determine Risk of Pathogen Transmission. PLoS ONE 8, e58656 (2013).

16. Sikulu, M. T. et al. Mass spectrometry identification of age-associated proteins from the malaria mosquitoes Anopheles gambiae s.s. and Anopheles stephensi. Data in Brief 4, 461–467 (2015).

17. Sikulu-Lord, M. T., Devine, G. J., Hugo, L. E. & Dowell, F. E. First report on the application of near-infrared spectroscopy to predict the age of Aedes albopictus Skuse. Sci Rep 8, 9590 (2018).

18. Ong, O. T. W. et al. Ability of near-infrared spectroscopy and chemometrics to predict the age of mosquitoes reared under different conditions. Parasites Vectors 13, 160 (2020).

19. Gao, Z. et al. Accurate age-grading of field-aged mosquitoes reared under ambient conditions using surface-enhanced Raman spectroscopy and artificial neural networks. Journal of Medical Entomology 60, (2023).

20. Wang, D. et al. Quantitative age grading of mosquitoes using surface-enhanced Raman spectroscopy. Analytical Science Advances 3, 47–53 (2022).

21. Krajacich, B. J. et al. Analysis of near infrared spectra for age-grading of wild populations of Anopheles gambiae. Parasites Vectors 10, 552 (2017).

22. Somé, B. M. et al. Adapting field-mosquito collection techniques in a perspective of near-infrared spectroscopy implementation. Parasites Vectors 15, 338 (2022).

23. Siria, D. J. et al. Rapid age-grading and species identification of natural mosquitoes for malaria surveillance. Nat Commun 13, 1501 (2022).

24. Mwanga, E. P. et al. Using transfer learning and dimensionality reduction techniques to improve generalisability of machine-learning predictions of mosquito ages from mid-infrared spectra. BMC Bioinformatics 24, 11 (2023).

25. González Jiménez, M., et al. Prediction of mosquito species and population age structure using mid-infrared spectroscopy and supervised machine learning [version 3; peer review: 2 approved]. Wellcome Open Research 4, (2019).

26. Johnson, B. J., Hugo, L. E., Churcher, T. S., Ong, O. T. W. & Devine, G. J. Mosquito Age Grading and Vector-Control Programmes. Trends in Parasitology 36, 39–51 (2020).

27. Gray, L. et al. Back to the Future: Quantifying Wing Wear as a Method to Measure Mosquito Age. The American Journal of Tropical Medicine and Hygiene 107, 689– 700 (2022).

28. Somé, A. F. et al. Safety and efficacy of repeat ivermectin mass drug administrations for malaria control (RIMDAMAL II): a phase 3, double-blind, placebo-controlled, cluster-randomised, parallel-group trial. The Lancet Infectious Diseases S1473309924007515 (2025) doi:10.1016/S1473-3099(24)00751-5.

29. Hugo, L. E., Quick-Miles, S., Kay, B. H. & Ryan, P. A. Evaluations of Mosquito Age Grading Techniques Based on Morphological Changes. JOURNAL OF MEDICAL ENTOMOLOGY 45, 17 (2008).

30. Anagonou, R. et al. Application of Polovodova’s method for the determination of physiological age and relationship between the level of parity and infectivity of Plasmodium falciparum in Anopheles gambiae s.s, south-eastern Benin. Parasit Vectors 8, 117 (2015).

31. Kirstein, O. D. et al. Targeted indoor residual insecticide applications shift Aedes aegypti age structure and arbovirus transmission potential. Sci Rep 13, 21271 (2023).

32. Christophers, S. The development of the egg follicle in anophelines. Paludism 1911, 73–88 (1911).

33. Bayili, K. et al. Evaluation of efficacy of Interceptor® G2, a long-lasting insecticide net coated with a mixture of chlorfenapyr and alpha-cypermethrin, against pyrethroid resistant Anopheles gambiae s.l. in Burkina Faso. Malar J 16, 190 (2017).

34. Mbewe, N. J. et al. Efficacy of bednets with dual insecticide-treated netting (Interceptor® G2) on side and roof panels against Anopheles arabiensis in north-eastern Tanzania. Parasites Vectors 15, 326 (2022).

35. N’Guessan, R., Odjo, A., Ngufor, C., Malone, D. & Rowland, M. A Chlorfenapyr Mixture Net Interceptor® G2 Shows High Efficacy and Wash Durability against Resistant Mosquitoes in West Africa. PLoS ONE 11, e0165925 (2016).

36. Tungu, P. K., Michael, E., Sudi, W., Kisinza, W. W. & Rowland, M. Efficacy of interceptor® G2, a long-lasting insecticide mixture net treated with chlorfenapyr and alpha-cypermethrin against Anopheles funestus: experimental hut trials in north-eastern Tanzania. Malar J 20, 180 (2021).

37. Lado, P. et al. Changing species dynamics and species-specific associations observed between Anopheles and Plasmodium genera in Diebougou health district, southwest Burkina Faso. Preprint at 10.1101/2024.10.09.24315124 (2024).

38. Slater, H. C. et al. Ivermectin as a novel complementary malaria control tool to reduce incidence and prevalence: a modelling study. The Lancet Infectious Diseases 20, 498–508 (2020).

